# SARS-CoV-2 spike conformation determines plasma neutralizing activity

**DOI:** 10.1101/2021.12.19.473391

**Authors:** John E. Bowen, Alexandra C. Walls, Anshu Joshi, Kaitlin R. Sprouse, Cameron Stewart, M. Alejandra Tortorici, Nicholas M. Franko, Jennifer K. Logue, Ignacio G. Mazzitelli, Sasha W Tiles, Kumail Ahmed, Asefa Shariq, Gyorgy Snell, Najeeha Talat Iqbal, Jorge Geffner, Alessandra Bandera, Andrea Gori, Renata Grifantini, Helen Y. Chu, Wesley C. Van Voorhis, Davide Corti, David Veesler

## Abstract

Numerous safe and effective COVID-19 vaccines have been developed that utilize various delivery technologies and engineering strategies. The influence of the SARS-CoV-2 spike (S) glycoprotein conformation on antibody responses induced by vaccination or infection in humans remains unknown. To address this question, we compared plasma antibodies elicited by six globally-distributed vaccines or infection and observed markedly higher binding titers for vaccines encoding a prefusion-stabilized S relative to other groups. Prefusion S binding titers positively correlated with plasma neutralizing activity, indicating that physical stabilization of the prefusion conformation enhances protection against SARS-CoV-2. We show that almost all plasma neutralizing activity is directed to prefusion S, in particular the S_1_ subunit, and that variant cross-neutralization is mediated solely by RBD-specific antibodies. Our data provide a quantitative framework for guiding future S engineering efforts to develop vaccines with higher resilience to the emergence of variants and longer durability than current technologies.

---

The SARS-CoV-2 spike (S) glycoprotein promotes viral entry into host cells and is the main target of neutralizing antibodies(*1, 2*). S comprises two functional subunits, designated S_1_ and S_2_, that interact non-covalently after furin cleavage during synthesis (*1, 3, 4*). The receptor-binding domain (RBD), which engages the ACE2 receptor (*1, 3, 5, 6*), and the N-terminal domain (NTD) that recognizes attachment factors (*7*–*9*) are components of the S_1_ subunit. The S_2_ subunit contains the fusion machinery and undergoes large-scale conformational changes to drive fusion of the virus and host membranes to initiate infection (*10, 11*). Antibodies that bind to specific sites on the RBD (*12*–*19*), the NTD (*20*–*23*), or the fusion machinery(*24*–*28*) neutralize SARS-CoV-2 and serum neutralizing antibody titers are a correlate of protection against SARS-CoV-2 (*29*–*34*).

As of December 2021, more than 7.8 billion COVID-19 vaccine doses have been administered from one of three different platforms: mRNA formulated with lipid nanoparticles, viral-vectored gene delivery or inactivated virus. Moderna/NIAID mRNA-1273 and Pfizer/BioNTech BNT162b2 were conceived as as two-dose vaccines based on an mRNA encoding the full-length prefusion-stabilized ‘2P’ S glycoprotein encapsulated in a lipid nanoparticle (*35*–*37*). AstraZeneca/Oxford AZD1222, Gamaleya Research Institute Sputnik V, and Janssen Ad26.COV2.S are replication-defective adenoviral-vectored vaccines encoding for the full-length S glycoprotein. Only Ad26.COV2.S encodes for a prefusion-stabilized S with the ‘2P’ mutations and removed furin cleavage site (*38*) whereas the other two vaccines lack these modifications. The adenoviral vectors used are chimpanzee AdY25 for AZD1222 (*39*) and Ad26 (prime)/Ad5 (boost) for Sputnik V (*40*), both vaccines initially employing two doses, and Ad26 for Ad26.COV2.S which originated as a single dose vaccine (*38*). Sinopharm BBIBP-CorV (*41*) is an alum-adjuvanted, β-propiolactone-inactivated SARS-CoV-2 viral vaccine which initially utilized a two dose regimen.

To understand the specificity of S-directed antibody responses elicited by vaccination, we evaluated plasma binding titers against the prefusion-stabilized SARS-CoV-2 S trimer, the NTD, the RBD, and the S_2_ subunit (fusion machinery) in the prefusion and postfusion states using enzyme-linked immunosorbent assay (ELISA). Our panel includes samples from individuals who received two doses of Moderna mRNA-1273, Pfizer/BioNTech BNT162b2, AstraZeneca AZD1222, Gamaleya Research Institute Sputnik V, or Sinopharm BBIBP-CorV, as well as individuals who received a single dose of Janssen Ad26.COV2.S. More than 3.5 billion doses of these vaccines have been administered worldwide as of December 2021. We benchmarked these samples against COVID-19 human convalescent plasma obtained before April 2021, likely resulting from exposure to a Washington-1-like isolate based on the date of symptom onset and the prevalence of this isolate in Washington State (*42*). Prefusion S binding titers were highest for individuals who had received two doses of mRNA-1273 or BNT162b2 (GMTs 1.8×10^4^ and 8.9×10^3^, respectively) and lowest for those who received a single dose of Ad26.COV2.S (GMT 2.1×10^2^) **(Fig. 1A, Fig. S1)**. The other two dose vaccines and SARS-CoV-2 infection resulted in intermediate prefusion S binding titers (GMT 1.0-1.4×10^3^) **(Fig. 1A, Fig. S1)**. Accordingly, the two mRNA vaccines induced greater magnitudes of RBD, NTD and prefusion S_2_ binding responses than all other groups **(Fig. 1A, Fig. S1)**.

**Figure 1.**
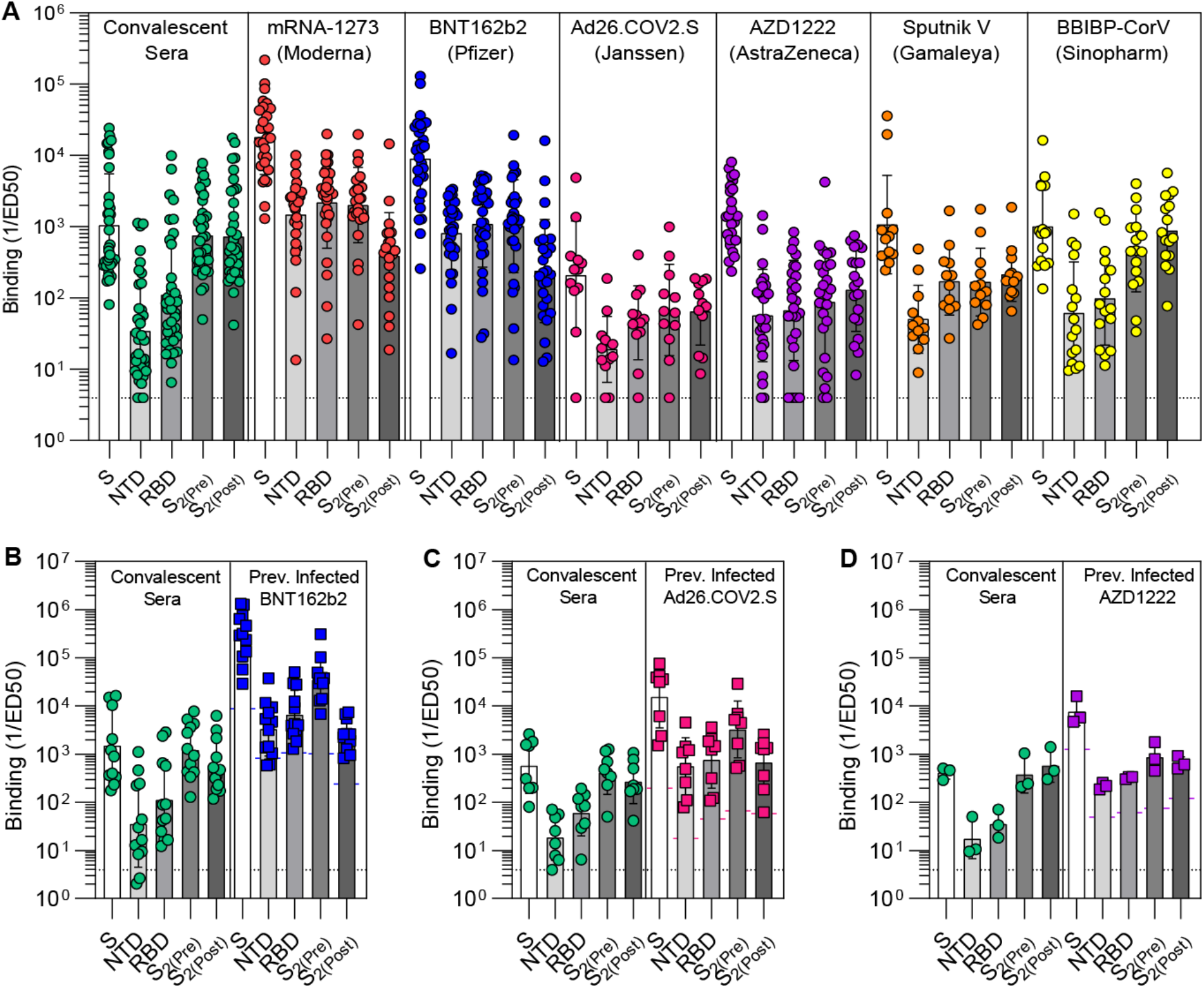
Prefusion-stabilization of SARS-CoV-2 S enhances S_1_ subunit antibody titers. (**A**) Antibody binding titers elicited by SARS-CoV-2 infection or vaccination to the prefusion S (S), the N-terminal domain (NTD), the receptor-binding domain (RBD), and the S_2_ subunit in the prefusion (S_2(Pre)_) and postfusion (S_2(Post)_) conformations, as measured by ELISA. (**B-D**) Antibody binding titers in matched cohorts of individuals previously infected with SARS-CoV-2 before and after vaccination with BNT162b2 (B), Ad26.COV2.S (C), or AZD1222 (D). Each point represents a single patient plasma sample, bars represent geometric means, and error bars represent geometric standard deviations. Protruding colored bars (B-D) mark the geometric mean of individuals that were not previously infected with SARS-CoV-2. Fit curves are shown Figure S1 and S2.

mRNA-1273 and BNT162b2 elicited polyclonal plasma antibodies with 5-fold greater prefusion to postfusion S_2_ binding titers **(Fig. 1A, Fig. S1)**, indicating preferential targeting of the prefusion state likely due to the ‘2P’ prefusion-stabilizing S mutations (*35*). Postfusion S_2_ binding titers for these two mRNA vaccines are likely accounted for by antibodies recognizing epitopes accessible in both conformations of the fusion machinery in ELISA assays (*28*) (**Fig. S4**). Conversely, natural infection or vaccination with AZD1222, Sputnik V or BBIBP-CorV, which do not contain prefusion-stabilizing S mutations, induced comparable prefusion and postfusion S_2_ binding titers **(Fig. 1A, Fig. S1)**. Furthermore, SARS-CoV-2 infection and BBIBP-CorV vaccination stood out due to their markedly low RBD-and NTD-specific, relative to postfusion S_2_-directed, antibody titers. These data point to a reduction of S_1_-directed antibodies relative to postfusion S_2_-targeting antibodies in these latter two groups likely due to S_1_ shedding and S_2_ refolding to the postfusion conformation at the surface of authentic virions or infected cells (*43*– *45*), or as a result of the β-propiolactone inactivation procedure utilized for producing BBIBP-CorV (*46*).

Previous studies demonstrated that administration of a single dose of vaccine to individuals previously infected with SARS-CoV-2 elicits high antibody binding and neutralizing titers (*47*–*50*). We therefore set out to assess and compare how binding responses are affected upon vaccination of previously infected subjects with two doses of BNT162b2, one dose of Ad26.COV2.S or two doses of AZD1222. Although vaccination led to a marked enhancement of antibody binding responses against all antigens tested, different vaccines led to distinct specific magnitudes of boosting. Comparison of post-vaccination to pre-vaccination prefusion S binding titers revealed increases of 2 orders of magnitude for BNT162b2 (184x) and one order of magnitude for Ad26.COV2.S (27x) and AZD1222 (19x) **(Fig. 1B-D, Fig. S2)**. BNT162b2 and Ad26.COV2.S vaccination induced greater prefusion S_2_ relative to postfusion S_2_ binding titers whereas AZD1222 resulted in comparable responses against both conformational states of the fusion machinery. The absence of prefusion-stabilizing S mutations in the latter vaccine might explain this outcome due to the metastable nature of the S trimer which is prone to shedding the S_1_ subunit and refolding to form postfusion trimers (*10, 35, 43, 45, 51*).

To investigate the relationship between antibody binding titers and neutralization potency, we determined half-maximum inhibitory dilutions of the aforementioned plasma samples using a vesicular stomatitis virus (VSV) pseudotyped with the Wuhan-Hu-1 S glycoprotein harboring the D614G substitution (G614) and VeroE6 cells stably expressing TMPRSS2(*52*). As a direct reflection of prefusion S binding titers, mRNA-1273 and BNT162b2 vaccinees’ plasma exhibited the highest neutralization potencies (GMTs 1,246 and 760, respectively) whereas Ad26.COV2.S-elicited plasma neutralizing activity after a single dose was the weakest among all groups (GMT 22) **(Fig. 2A, Fig. S3A)**. Neutralizing antibody GMTs for previously exposed individuals reached 7,423 after BNT162b2 vaccination, 1,028 after Ad26.COV2.S vaccination, and 1,251 after AZD1222 vaccination **(Fig. 2A, Fig. S3A)**, a 1-2 order of magnitude rise over corresponding vaccinees who had not been previously exposed to SARS-CoV-2 (*47*–*50*).

**Figure 2.**
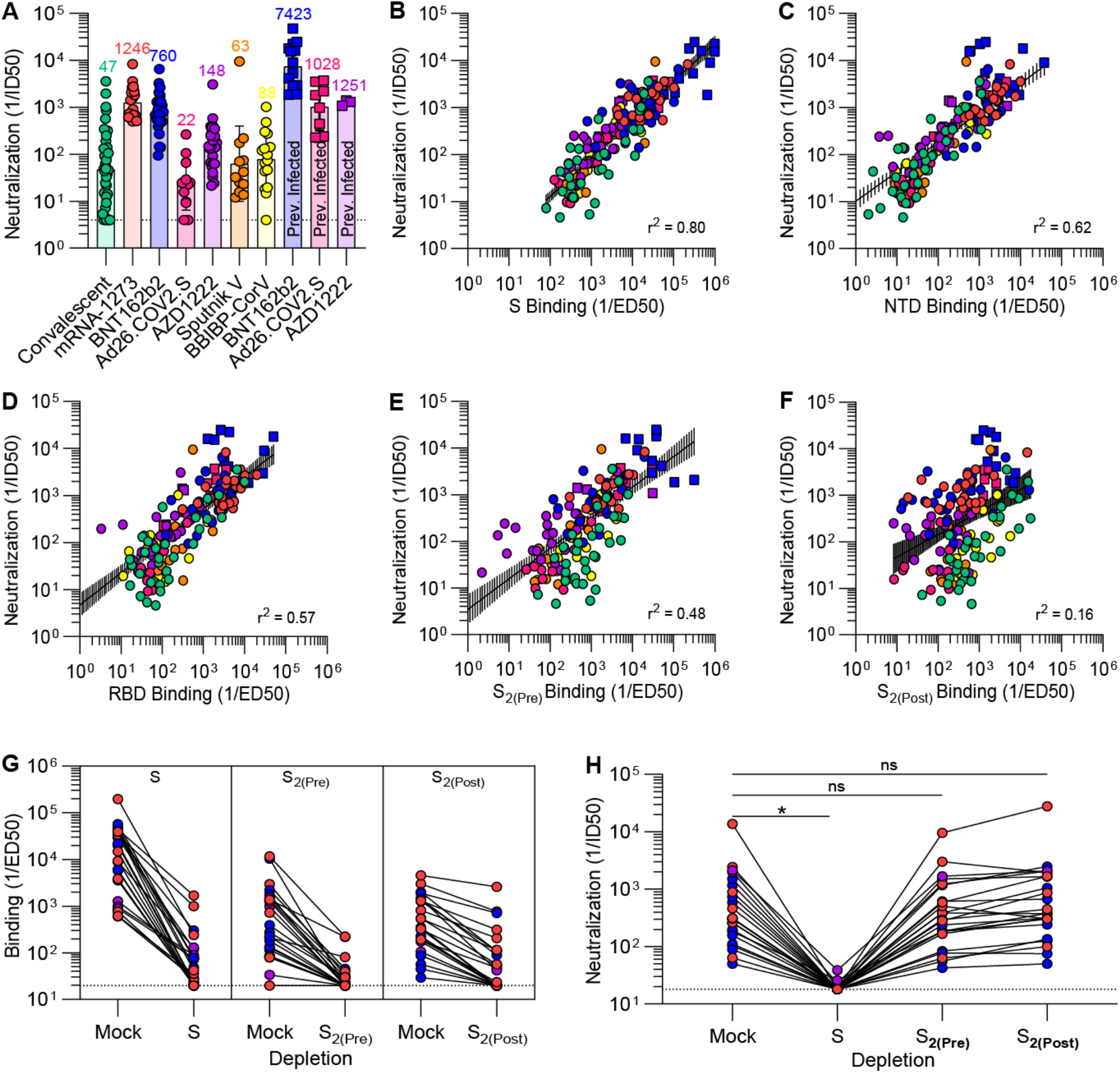
SARS-CoV-2 neutralization is determined by S_1_ subunit targeting antibodies. (**A**) SARS-CoV-2 S pseudotyped VSV neutralization titers elicited by natural infection or vaccination. The dotted line is the limit of detection, the colored bars are GMTs and the black error bars are geometric standard deviations. Fit curves are shown in Fig. S3. (**B-F**) Correlation between plasma neutralizing activity and prefusion S (B), N-terminal domain (C), receptor-binding domain (D), prefusion S_2_ (E), and postfusion S_2_ (F) binding titers shown with a linear regression fit to the log of neutralization and binding titers. The black shaded regions represent 95% confidence intervals. (**G-H**) Binding (G) and neutralization (H) titers resulting from depletion of polyclonal antibodies targeting prefusion S, prefusion S_2_, and postfusion S_2_. Fit curves are shown in Fig. S5 and S6. Statistical significance between groups of paired data were determined by Wilcoxon rank test and *P < 0.05.

We observed a strong positive correlation between in vitro plasma inhibitory activity and the magnitude of antibody responses against the prefusion-stabilized S trimer for all vaccines evaluated and for infection-elicited polyclonal antibodies **(Fig. 2B)**. Neutralizing antibody titers were also correlated with NTD- and RBD-specific binding titers **(Fig. 2C-D)**, in line with these two domains being the main targets of neutralizing antibodies upon infection or vaccination (*15, 20, 47, 53*–*55*). The rapid accumulation of residue mutations in the SARS-CoV-2 S_1_ subunit throughout the COVID-19 pandemic might therefore reflect (at least in part) the selective pressure exerted by host neutralizing antibodies (*56*). Although prefusion S_2_ antibody titers had a relatively strong correlation with neutralization potency **(Fig. 2E)**, postfusion S_2_ responses did not **(Fig. 2F)**, indicating that antibodies targeting postfusion S are likely to have a limited impactt on reducing viral entry. Natural infection elicited the most heterogeneous humoral immune responses as defined by the widespread of prefusion S binding and associated neutralizing antibody titers compared to other groups **(Fig. 1A and Fig. 2A)**. Vaccination of previously infected individuals with a single dose of Ad26.COV2.S or two doses of AZD1222 elicited neutralizing antibody titers comparable or greater than two doses of mRNA-1273 or BNT162b2 in naive individuals, in line with previous reports (*47*–*50*). Collectively, these results emphasize the benefits of favoring the prefusion S conformation to maximize elicited neutralizing antibody titers as well as the higher quality of humoral immune response elicited by vaccination with most platforms compared to natural infection.

To obtain a quantitative understanding of the relationship between S conformation and plasma neutralizing activity, we depleted polyclonal antibodies using prefusion S, prefusion S_2_ or postfusion S_2_ from the plasma of vaccinees who had received mRNA-1273, BNT162b2, or AZD1222. Binding titers against the respective antigens were reduced by ≥1.5 orders of magnitude as determined by ELISA, confirming effective antigen-specific antibody removal (**Fig.2G**). Depletion of prefusion S-targeting antibodies resulted in a ≥91% (mRNA-1273), ≥96% (BNT162b2) and ≥89% (AZD1222) reduction of neutralizing GMT whereas depletion using prefusion or postfusion S_2_ only had a modest impact **(Fig. 2H)**. These results clearly demonstrate that virtually all plasma neutralizing activity targets the prefusion S conformation, which is reminiscent of findings made for the respiratory syncytial virus fusion (F) glycoprotein (*57, 58*). The coronavirus S glycoprotein, however, mediates both receptor-binding and membrane fusion whereas respiratory syncytial virus F solely promotes membrane fusion. Our data indicate that it is S_1_ subunit shedding, rather than S_2_ conformational changes, that leads to a marked loss of potency due to the fact that most neutralizing antibodies target the RBD (*15, 53, 59*) and to a lesser extent the NTD (*20, 21, 23*).

To evaluate the relative contribution of the RBD and the NTD to cross-neutralizing activity against SARS-CoV-2 variants, we depleted plasma samples of antibodies recognizing each of these two antigens using vaccinees who received two doses of either BNT162b2 (8 individuals) or mRNA-1273 (8 individuals) **(Fig. 3A)**. Plasma neutralizing activity was subsequently determined against VSV pseudotyped with the G614 S glycoprotein, the Delta (B.1.617.2) S, or the Omicron S (B.1.1.529) using VeroE6 cells stably expressing TMPRSS2 (*52, 55, 60, 61*). Whereas depletion of NTD-specific antibodies reduced neutralization GMT by up to 67% against G614 S VSV, it had no effect on inhibition of the two variants tested **(Fig. 3B-D)**. In contrast, depletion of RBD-specific antibodies completely abrogated variant cross-neutralization, suggesting that vaccine-elicited neutralization breadth was almost completely accounted for by antibodies targeting this domain **(Fig. 3B-D)**. These data concur with (i) the marked antigenic variation of the SARS-CoV-2 NTD among variants and sarbecoviruses, which is associated with a narrow specificity of NTD neutralizing antibodies (*20, 23, 54, 61*–*63*), and (ii) the description of multiple broadly neutralizing antibodies recognizing distinct RBD antigenic sites (*12*–*14, 16, 18, 64*–*67*).

**Figure 3.**
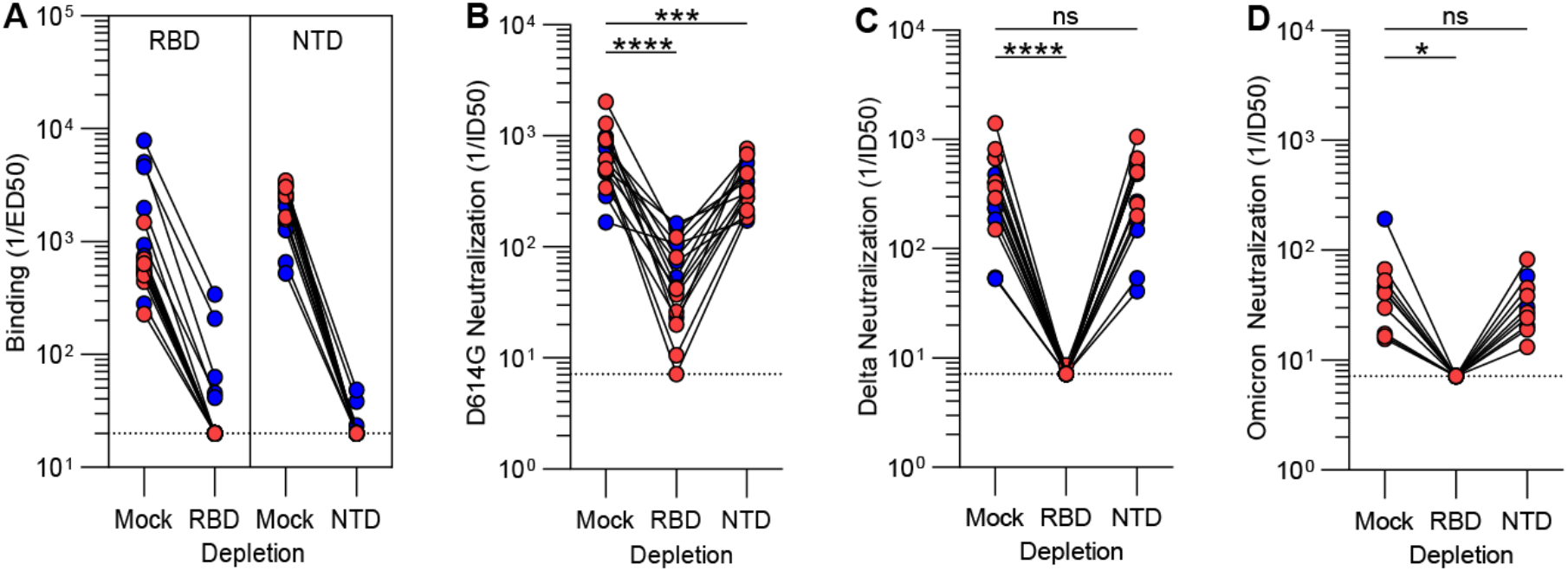
Broad neutralization of SARS-CoV-2 variants is mediated by RBD-specific antibodies. (**A-B**) Plasma binding titers resulting from mock, Wuhan-Hu-1 RBD (left) and NTD (right) depletion of polyclonal antibodies. (**B-D**) Plasma neutralizing activity against G614 S VSV (B), Delta S VSV (C) and Omicron S VSV (D) after mock, Wuhan-Hu-1 RBD or NTD depletion of polyclonal antibodies. Mock consists of depletion carried out with beads without immobilized antigen. Statistical significance between groups of paired data were determined by Wilcoxon rank test and *P < 0.05, ***P < 0.001, ****P < 0.0001.

The discovery that most neutralizing activity in subjects infected with respiratory syncytial virus targets prefusion F led to subsequent stabilization of this conformational state through protein engineering and yielded lead vaccine candidates against this pathogen (*57, 58, 68, 69*). We demonstrate here that prefusion SARS-CoV-2 S binding titers correlate with plasma neutralizing activity largely due to targeting of the S_1_ subunit, which comprises antigenic sites recognized by most neutralizing antibodies and is shed upon refolding. Targeting of the prefusion S_2_ subunit conformation, as opposed to the postfusion state, makes only a small contribution to overall neutralizing activity due to the low frequency and potency of fusion machinery-specific neutralizing antibodies (*24, 28, 70*). The data presented here in humans concur with mouse and non-human primate immunogenicity studies showing that prefusion-stabilized ‘2P’ S glycoproteins elicit greater neutralizing antibody titers than non-stabilized S trimers (*35, 38, 71*). These outcomes are likely resulting from the metastability of prefusion S and suggest that utilization of next-generation S immunogens with additional perfusion-stabilizing mutations (e.g. ‘HexaPro S’ (*72*) or ‘VLFIP’ S (*73*)) could lead to vaccines eliciting even greater neutralizing antibody titers and in turn enhanced durability and resilience to SARS-CoV-2 variants. The identification of the RBD as the sole target of vaccine-elicited polyclonal antibodies broadly neutralizing SARS-CoV-2 variants is reminiscent of recent reports describing broadly neutralizing sarbecovirus monoclonal antibodies isolated from infected subjects (*12*–*14, 16, 18, 64*–*67*) and the rapid accumulations of mutations in the NTD (*20, 54, 61*–*63*). These findings motivate the clinical development of RBD-based vaccines against SARS-CoV-2 (*29, 74*–*77*) and sarbecoviruses (*78*) for future pandemic preparedness.

## Author contributions

J.E.B., A.C.W., M.A.T. and D.V. conceived the project and designed experiments. J.E.B. performed all experiments. J.E.B. and K.S. produced pseudotyped viruses. J.E.B., A.J., and C.S. expressed and purified proteins. I.G.M., N.T.I., G.S., J.G., A.B., A.G., R.G., H.Y.C, W.C.V.V., and D.C. provided unique reagents. D.V. supervised the project and obtained funding. J.E.B. and D.V. analyzed the data and wrote the manuscript with input from all authors.

## Acknowledgements

We thank Hideki Tani (University of Toyama) for providing the reagents necessary for preparing VSV pseudotyped viruses. This study was supported by the National Institute of Allergy and Infectious Diseases (DP1AI158186 and HHSN272201700059C to D.V., U01 AI151698 to W.C.V.V.), the National Institute of General Medical Sciences (R01GM120553 to D.V.), a Pew Biomedical Scholars Award (D.V.), an Investigators in the Pathogenesis of Infectious Disease Awards from the Burroughs Wellcome Fund (D.V.), Fast Grants (D.V.), the Bill & Melinda Gates Foundation (OPP1156262 to D.V.), the University of Washington Arnold and Mabel Beckman cryoEM center and the National Institute of Health grant S10OD032290 (to D.V.) and grant U01 AI151698 for the United World Antiviral Research Network (UWARN) as part of the Centers for Research in Emerging Infectious Diseases (CREID) Network. D.V. is an Investigator of the Howard Hughes Medical Institute.

## Supplemental Material

**Figure S1.**
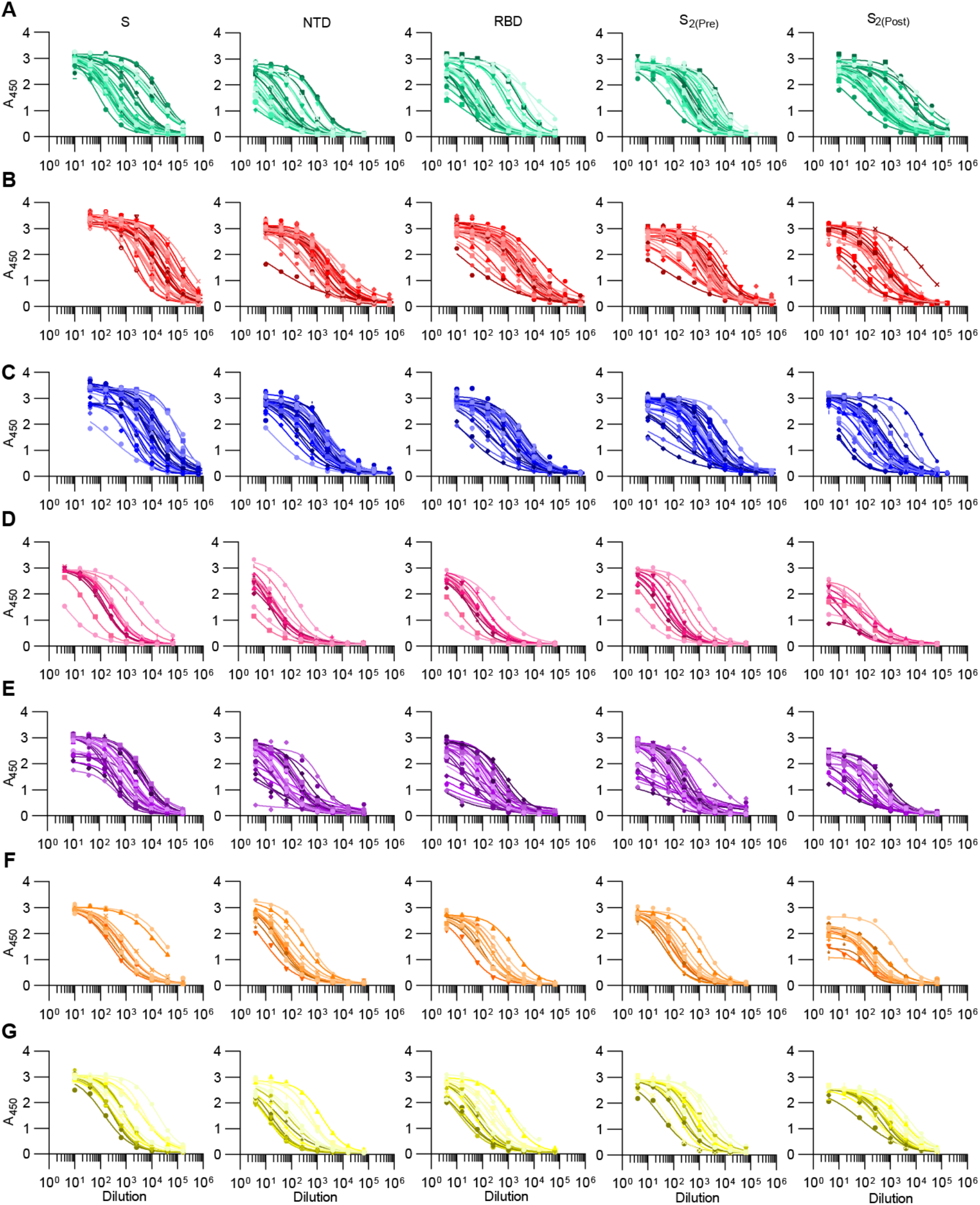
Fit ELISA curves of plasma antibody binding to the SARS-CoV-2 S, the N-terminal domain, the receptor-binding domain, and the S_2_ subunit in the prefusion and postfusion conformations (left to right). Samples were collected from individuals after SARS-CoV-2 infection (A) or vaccination with mRNA-1273 (B), BNT162b2 (C), Ad26.COV2.S (D), AZD1222 (E), Sputnik V (F), and BBIBP-CorV (E). All vaccines consisted of two doses besides Ad26.COV2.S which consisted of only one.

**Figure S2.**
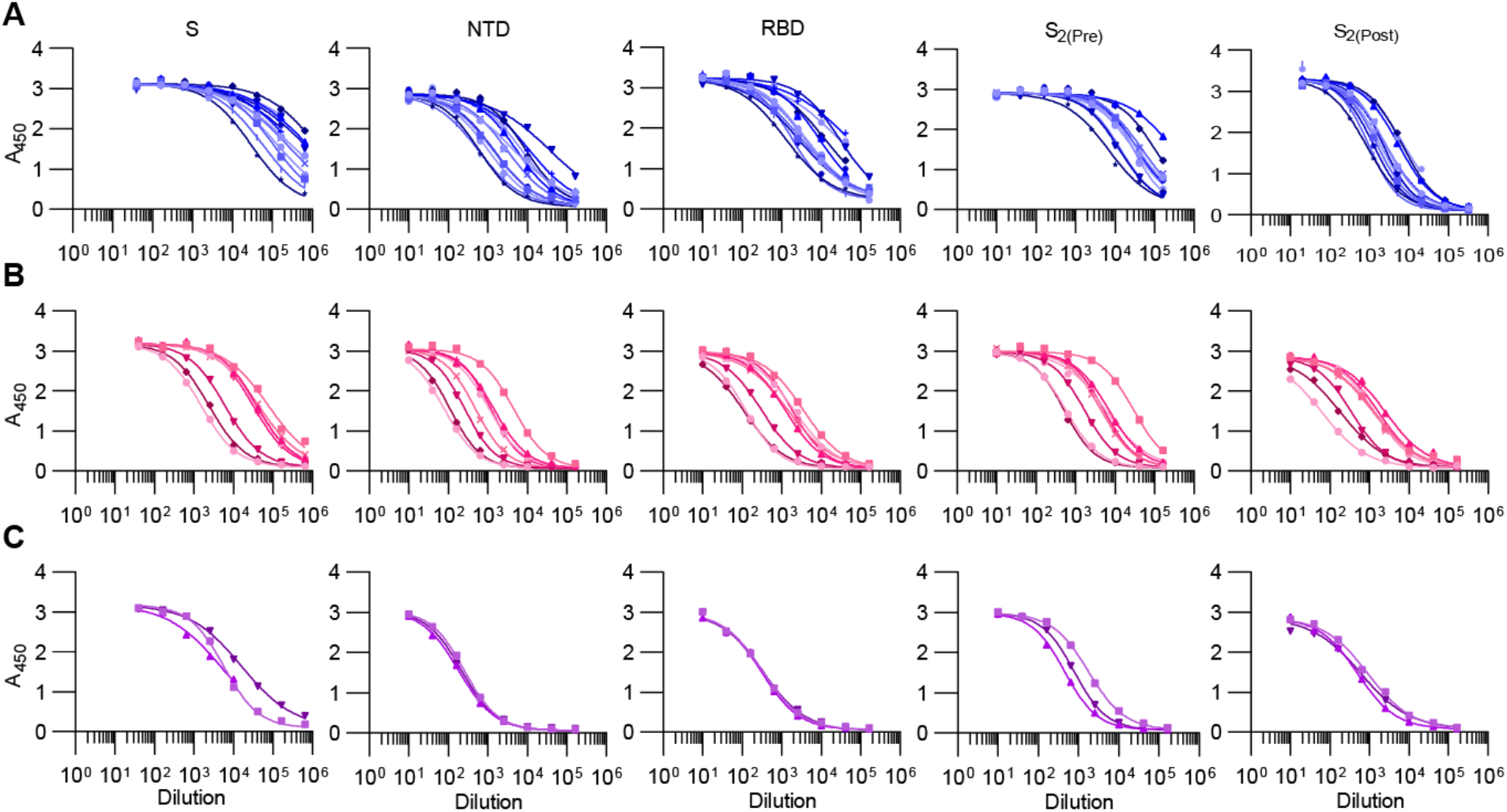
Fit ELISA curves of plasma antibody binding to the SARS-CoV-2 S, the N-terminal domain, the receptor-binding domain, and the S_2_ subunit in the prefusion and postfusion conformations (left to right). Samples were collected from individuals after SARS-CoV-2 infection and vaccination with BNT162b2 (A), Ad26.COV2.S (B), and AZD1222 (C). BNT162b2 and AZD1222 consisted of two doses whereas Ad26.COV2.S consisted of only one.

**Figure S3.**
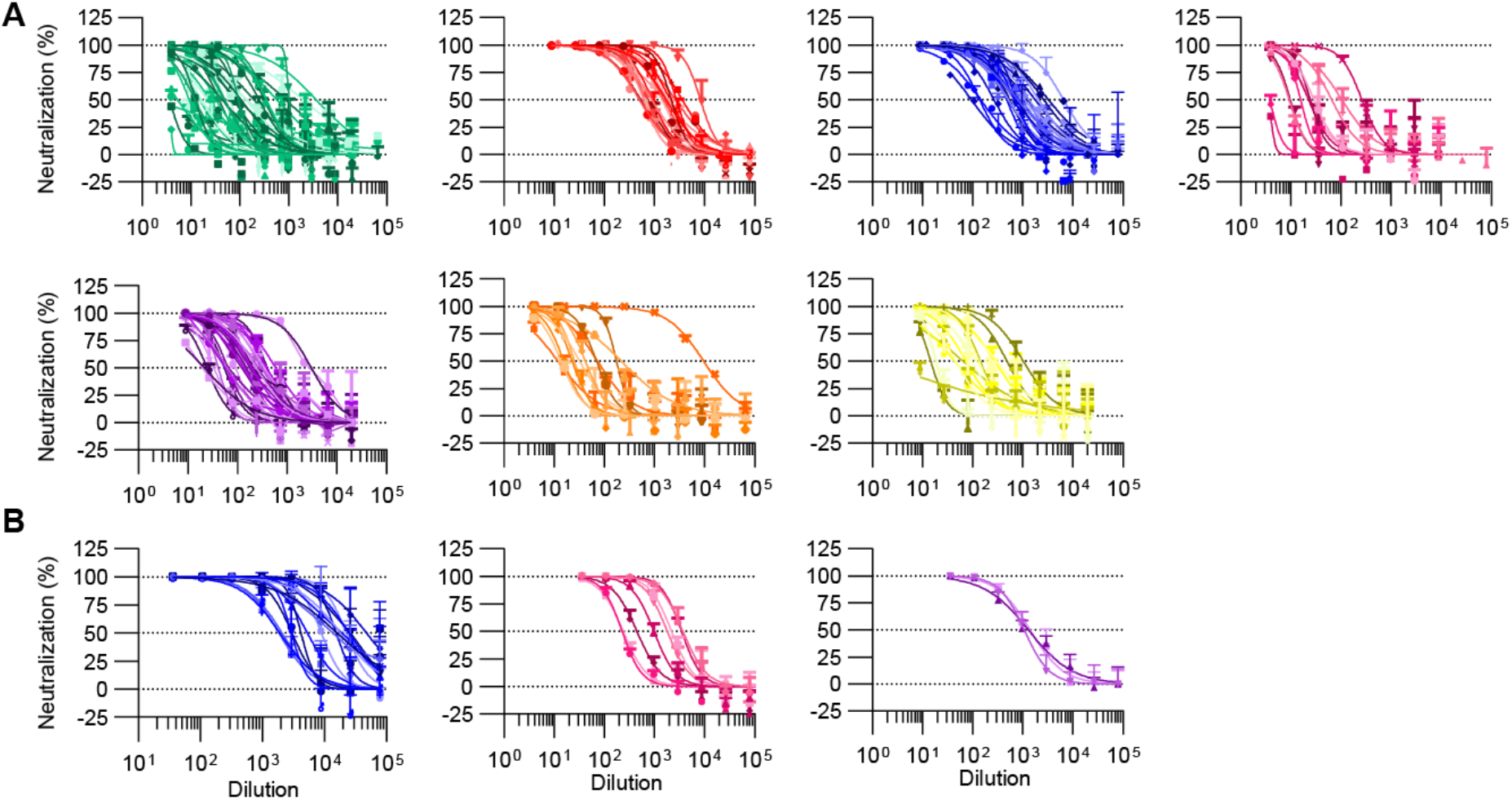
Normalized neutralization curves using SARS-CoV-2 S D614G VSV pseudovirus on VeroE6-TMPRSS2 cells. (**A**) Samples were collected from individuals after SARS-CoV-2 infection (green) or vaccination with mRNA-1273 (red), BNT162b2 (blue), Ad26.COV2.S (pink), AZD1222 (purple), Sputnik V (orange), or BBIBP-CorV (yellow). All vaccines consisted of two doses besides Ad26.COV2.S which consisted of only one. (**B**) Samples were also collected from individuals after SARS-CoV-2 infection and vaccination with BNT162b2 (blue), Ad26.COV2.S (pink), and AZD1222 (purple). BNT162b2 and AZD1222 consisted of two doses whereas Ad26.COV2.S consisted of only one.

**Figure S4.**
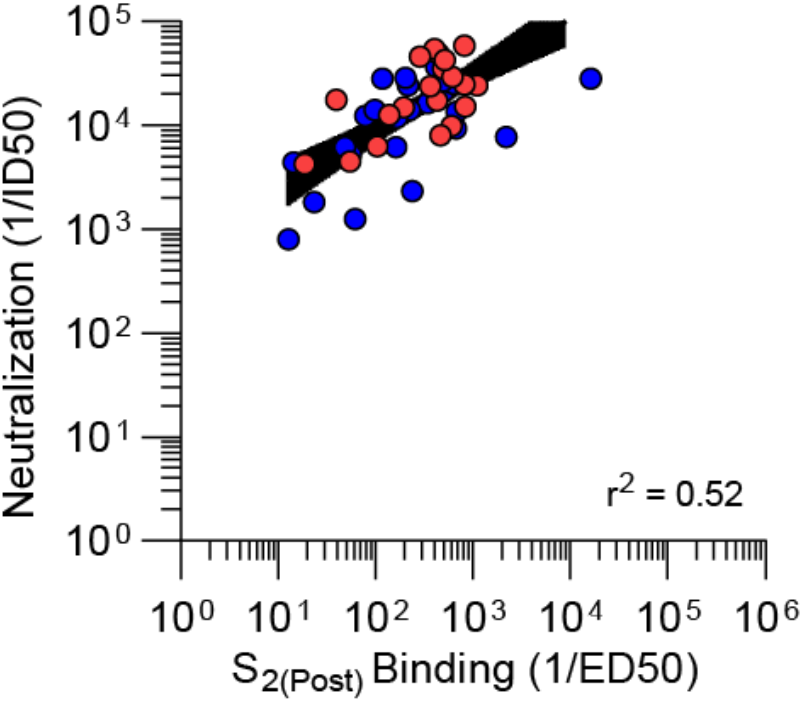
Correlation between SARS-CoV-2 neutralization titers and binding titers to the S_2_ subunit in the postfusion conformation of individuals vaccinated with two doses of mRNA-1273 (red) or BNT162b2 (blue).

**Figure S5.**
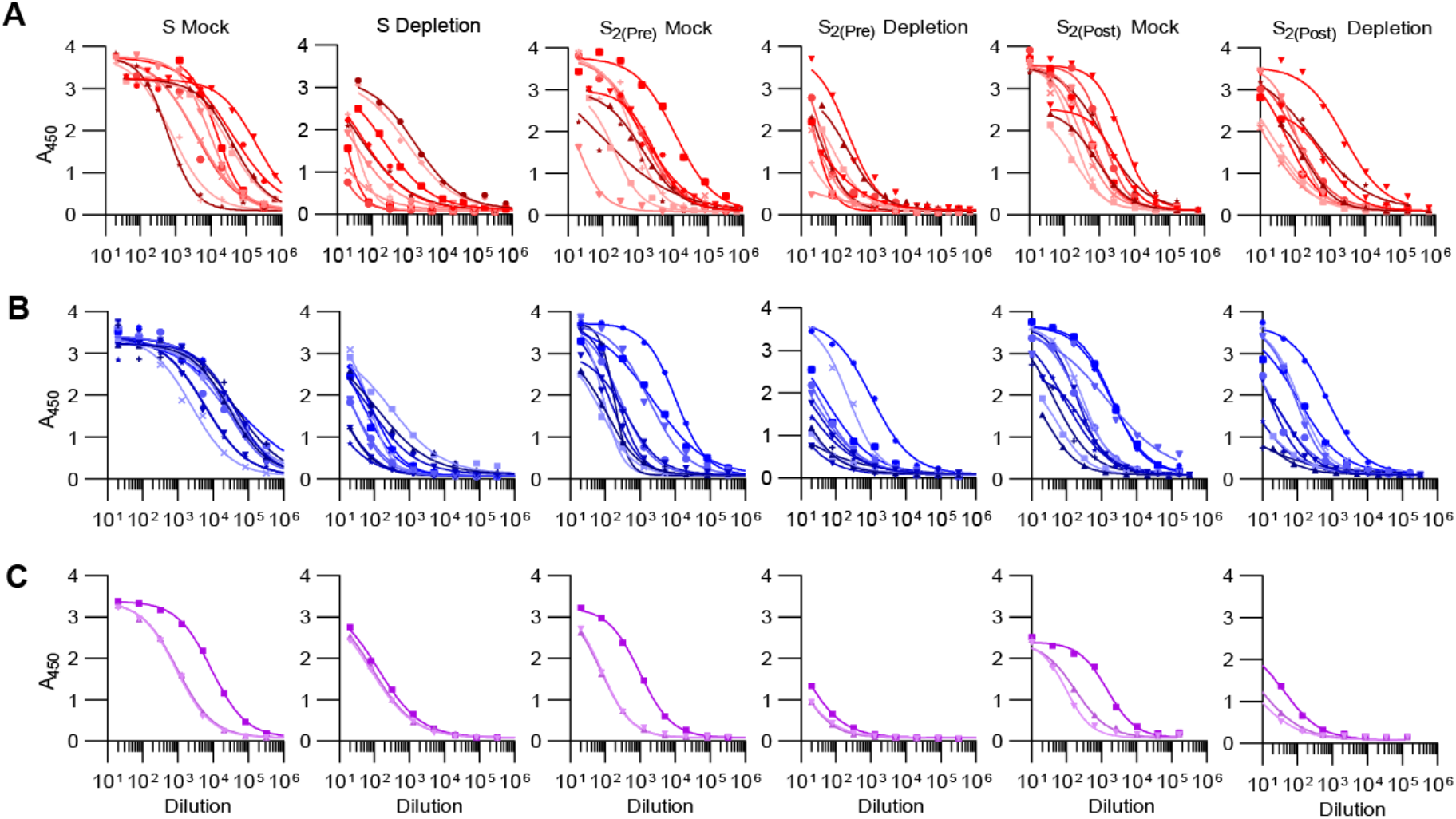
Fit ELISA curves of plasma antibody binding to SARS-CoV-2 S, S_2_ in the prefusion conformation, or S_2_ in the postfusion conformation following mock or protein depletion. Samples were collected from individuals vaccinated with two doses of mRNA-1273 (C), BNT162b2 (B), or AZD1222 (C).

**Figure S6.**
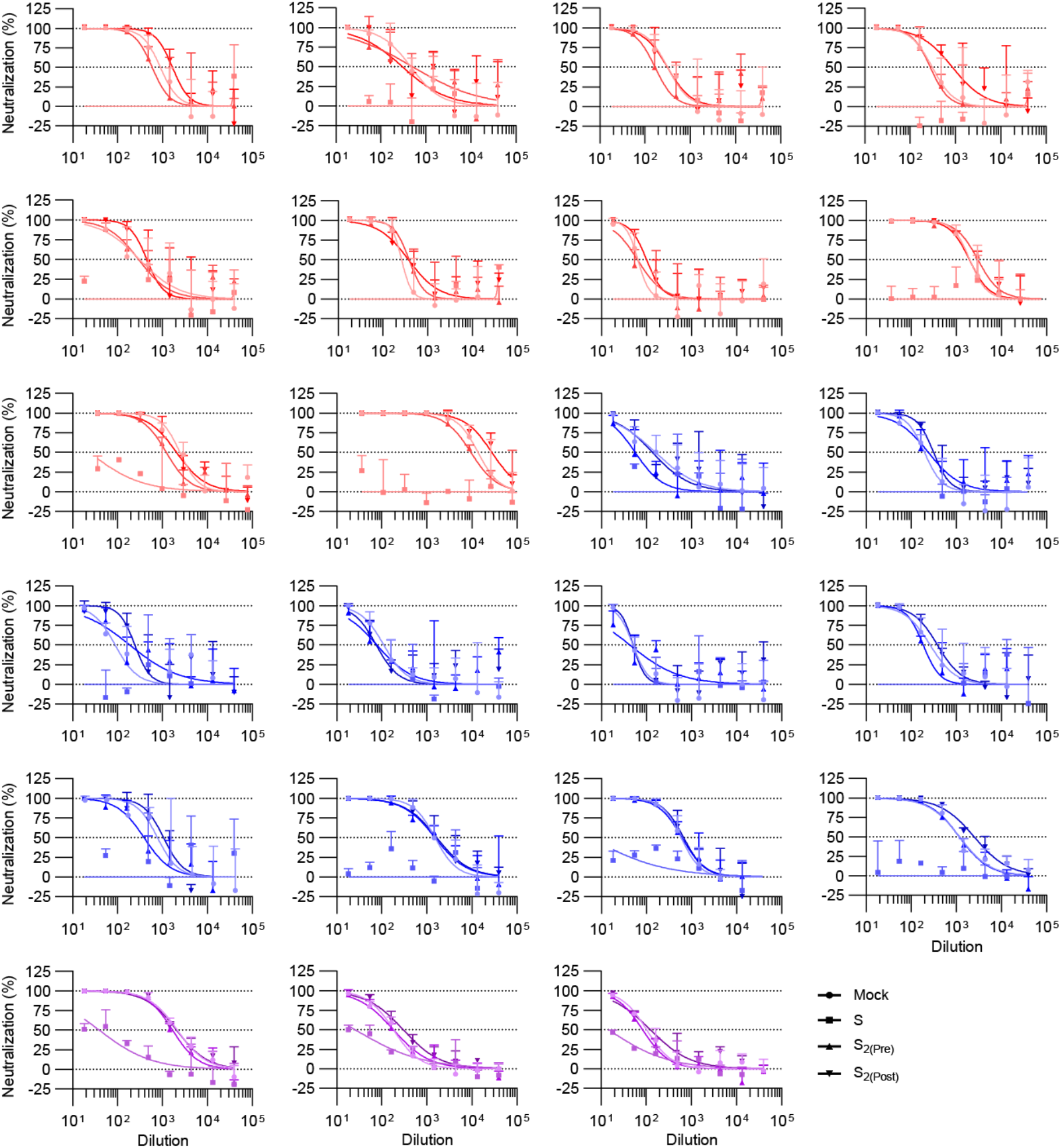
Normalized neutralization curves using SARS-CoV-2 S D614G VSV pseudovirus on VeroE6-TMPRSS2 cells following mock depletion or depletion with S, S_2_ in the prefusion conformation, or S_2_ in the postfusion conformation.

**Figure S7.**
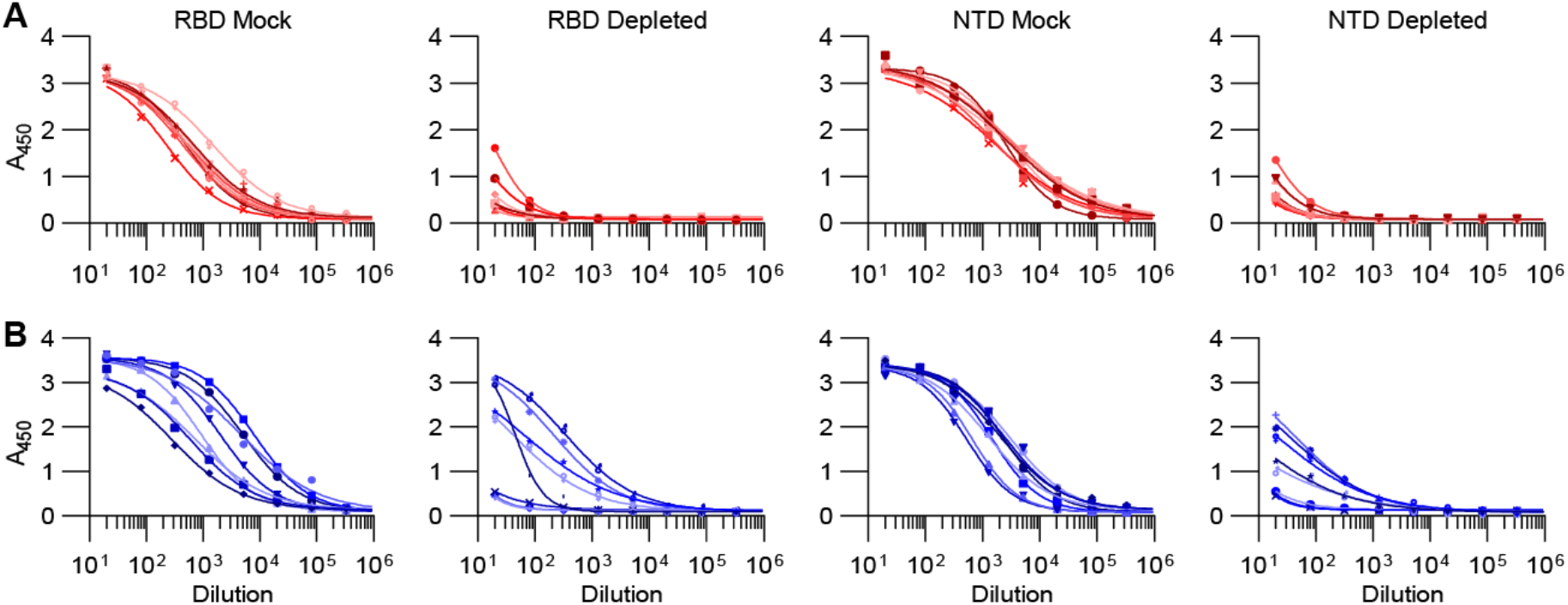
Fit ELISA curves of plasma antibody binding to the RBD or NTD following mock or protein depletion. Samples were collected from individuals vaccinated with two doses of mRNA-1273 (A) or BNT162b12 (B).

**Figure S8.**
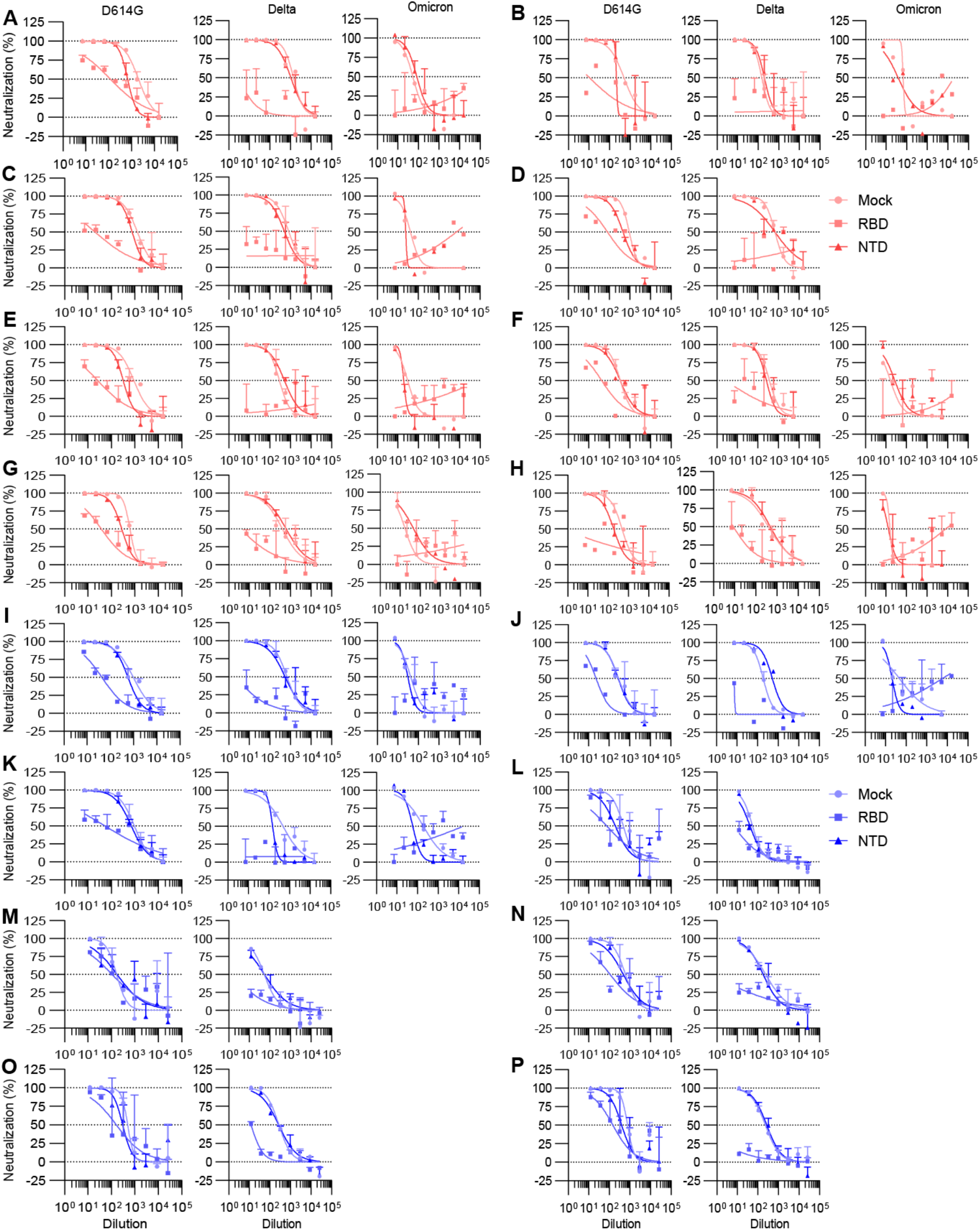
Normalized neutralization curves using SARS-CoV-2 D614G S, Delta S, or Omicron S VSV pseudoviruses on VeroE6-TMPRSS2 cells following mock depletion or depletion with Wuhan-Hu-1 RBD or NTD. Samples were collected from individuals vaccinated with two doses of mRNA-1273 (A-H) or BNT162b12 (I-P).

**Table S1.**
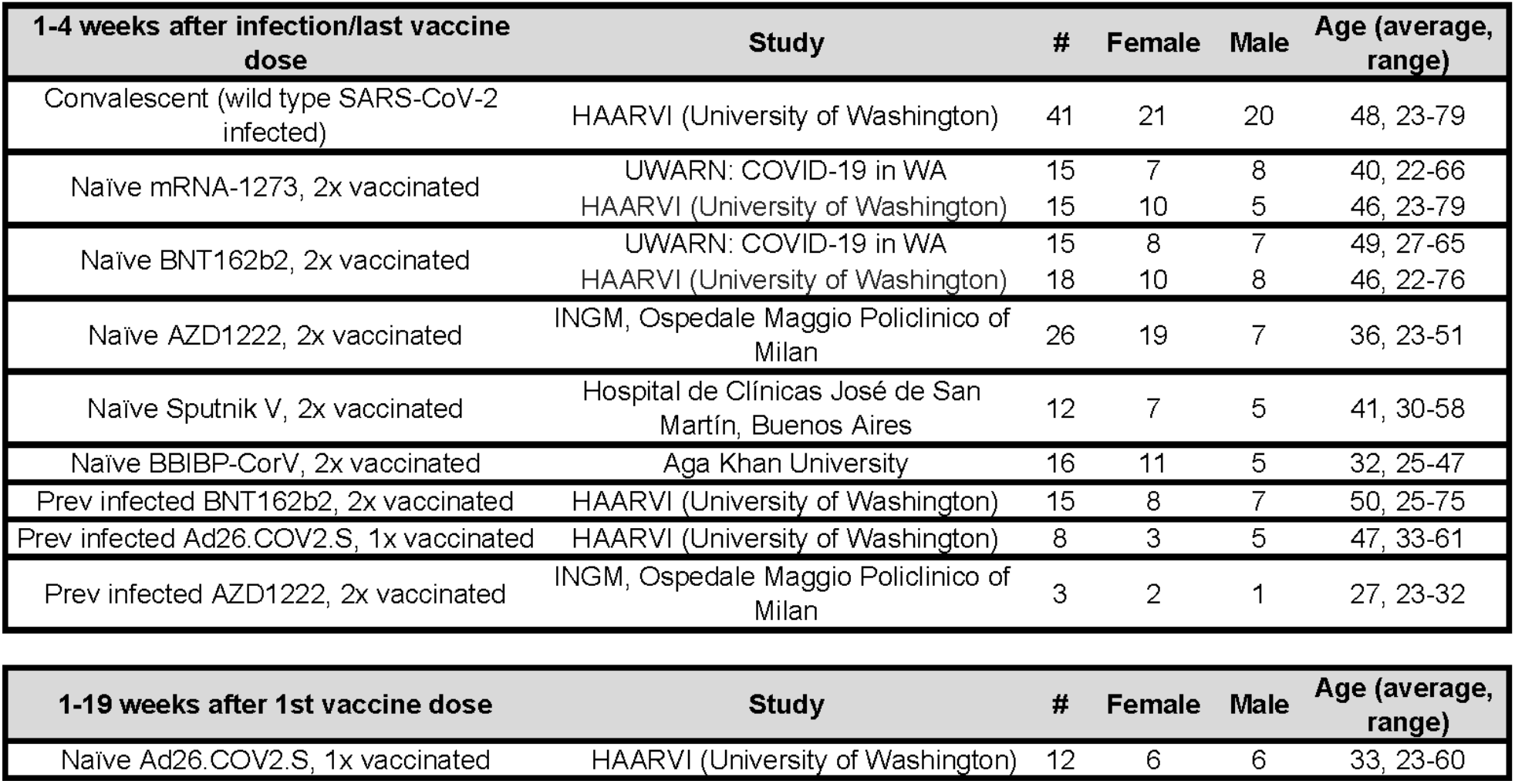
Demographics data of enrolled plasma donors.

## Methods

### Cell lines

Cell lines used in this study were obtained from ThermoFisher Scientific (HEK293T and Expi293F) or were kindly gifted by Florian Lempp (Vero-TMPRSS2 cells(*52*)). None of the cell lines used were authenticated or tested for mycoplasma contamination.

### Sample donors

Convalescent plasma, Ad26.COV2.S, and some BNT162b2 samples were obtained from the HAARVI study approved by the University of Washington Human Subjects Division Institutional Review Board (STUDY00000959). mRNA-1273 and BNT162b2 samples were obtained from individuals enrolled in the UWARN: COVID-19 in WA study approved by the University of Washington Human Subjects Division Institutional Review Board (STUDY00010350). AZD1222 samples were obtained from INGM, Ospedale Maggio Policlinico of Milan and approved by the local review board Study Polimmune. Sputnik V samples were obtained from healthcare workers at the hospital de Clínicas “José de San Martín”, Buenos Aires, Argentina. BBIBP-CorV samples were obtained from Aga Khan University, Karachi, Pakistan. Demographic data for these individuals are summarized in Table 1.

### Plasmid construction

The SARS-CoV-2 S-Hexapro is as previously described (Hsieh et al., 2020) and placed into CMVR with an octa-his tag. The SARS-CoV-2 N-terminal domain:SARS-CoV-2 NTD (residues 14-307) with a C-terminal 8XHis-tag was sub-cloned in pCMV as previously described (McCallum et al., 2020).The SARS-CoV-2-RBD-Avi construct was synthesized by GenScript into pcDNA3.1-with an N-terminal mu-phosphatase signal peptide and a C-terminal octa-histidine tag, flexible linker, and avi tag (GHHHHHHHHGGSSGLNDIFEAQKIEWHE). The boundaries of the construct are N-_328_RFPN_331_ and _528_KKST_531_-C (Walls et al., 2020a). SARS-CoV-2 S-D614G (Lempp et al. 2021) has a mu-phosphatase signal peptide beginning at Q14, a mutated S1/S2 cleavage site (SGAR), ends at residue K1211 and is followed by a TEV cleavage site, fold-on trimerization motif, and an 8× His tag in the pCMV vector. The SARS-CoV-2 S_2_-pentapro has a mu-phsophatase signal peptide and begins at _697_MSLG_700_ with ⅚ of the proline stabilizing mutations as described in Hsieh et al. 2020. The proline at position 817 was not used in the pentapro construct. The construct ends at residue K1211 and is followed by a TEV cleavage site, fold-on trimerization motif, and an 8× His tag in the CMVR vector. SARS-CoV-2 G614 S (YP 009724390.1), and Delta (B.1.617.2) S genes were all placed into the HDM vector with a 21 residue C-terminal deletion, as previously described (*63, 78, 79*). The plasmids encoding the SARS-CoV-2 Omicron (B.1.1.529) S variant was generated by overlap PCR mutagenesis of the wildtype plasmid, pcDNA3.1(+)-spike-D19 (*80*).

### Protein expression and purification

Proteins were produced in Expi293F Cells (ThermoFisher Scientific) grown in suspension using Expi293 Expression Medium (ThermoFisher Scientific) at 37°C in a humidified 8% CO2 incubator rotating at 130 rpm. Cells grown to a density of 3 million cells per mL were transfected using the ExpiFectamine 293 Transfection Kit (ThermoFisher Scientific) and cultivated for 3-5 days. Proteins were purified from clarified supernatants using a nickel HisTrap HP affinity column (Cytiva) and washed with ten column volumes of 20 mM imidazole, 25 mM sodium phosphate pH 8.0, and 300 mM NaCl before elution on a gradient to 500 mM imidazole. To produce SARS-CoV-2 S in the postfusion state, SARS-CoV-2 S D614G was incubated with 1:1 w/w S2X58-Fab (*14*) and 10 ug/mL trypsin for one hour at 37C before size exclusion on a Superose 6 Increase column (Cytivia). Proteins were buffer exchanged into 20 mM sodium phosphate pH 8 and 100 mM NaCl and concentrated using centrifugal filters (Amicon Ultra) before being flash frozen.

### Enzyme-linked immunosorbent assay (ELISA)

30 µL of 3 ug/mL of SARS-CoV-2 S Hexapro, NTD, RBD, S_2(Pre)_, or S_2(Post)_ diluted in PBS were incubated on a 384-well Nunc Maxisorp plate (ThermoFisher 464718) for one hour at 37C. Plates were slapped dry before addition of 80 µL blocker Casein in PBS (ThermoFisher) and incubation for one hour at 37C. Plates were slapped dry and a 1:4 serial dilution of plasma in 30 µL TBST was added and incubated for one hour at 37C. Plates were slapped dry and washed 4x with TBST using a BioTek plate washer followed by addition of Invitrogen anti-Human IgG (ThermoFisher A18817) and one hour incubation at 37C. Plates were once again slapped dry and washed 4x with TBST before addition of room temperature TMB Microwell Peroxidase (Seracare 5120-0083). The reaction was quenched after 1-2 minutes with 1 N HCl and the A450 of each well was read using a BioTek plate reader. Prism (GraphPad) nonlinear regression with “Sigmoidal, 4PL, X is concentration” was used to determine the ED50 of each sample. The bottom was constrained to the minimum A450 per plate and top was constrained to the maximum A450 per plate, besides postfusion S binding for Ad26.COV2.S and Sputnik V where the top was constrained to less than the maximum A450 per plate due to wide variability in maximum A450.

### Pseudotyped VSV production

SARS-CoV-2 D614G, Delta (B.1.617.2), and Omicron (B.1.1.529) pseudotypes were prepared similarly as previously described (*63*). Briefly, HEK-293T cells seeded in poly-D-lysine coated 100 mm dishes at ∼75 % confluency were washed five times with Opti-MEM and transfected using 24 µg of the S glycoprotein plasmid with Lipofectamine 2000 (Life Technologies). After 5 h at 37°C, media supplemented with 20% FBS and 2% PenStrep was added. After 20 hours, cells were washed five times with DMEM and cells were transduced with VSVΔG-luc before a 2 h incubation at 37°C. Infected cells were then washed an additional five times with DMEM prior to adding media supplemented with anti-VSV-G antibody (I1-mouse hybridoma supernatant diluted 1:25, from CRL-2700, ATCC) to reduce parental background. After 18-24 h, the supernatant was harvested and clarified by low-speed centrifugation at 2,500 g for 10 min. The supernatant was then filtered (0.45 μm) and concentrated 10 times using a 30 kDa centrifugal concentrator (Amicon Ultra). The pseudotypes were then aliquoted and frozen at -80 °C.

### Pseudotyped VSV neutralization assay

To evaluate neutralization of D614G, Delta (B.1.617.2), and Omicron (B.1.1.529) pseudotypes by plasma of vaccinees or previously infected individuals, Vero-TMPRSS2 cells in DMEM supplemented with 10% FBS, 1% PenStrep, and 8 ug/mL puromycin were seeded at 60-70% confluency into white clear-bottom 96 well plates (Corning) and incubated at 37°C. The following day, a half-area 96-well plate (Greiner) was prepared with eight 3-fold serial plasma dilutions. An equal volume of DMEM with 1:25 pseudovirus and 1:25 anti-VSV-G antibody (I1-mouse hybridoma supernatant from CRL-2700, ATCC) was then added to the half-area plate. The mixture was incubated at room temperature for 20-30 minutes. Media was removed from the cells and 40 µL from each well (containing plasma and pseudovirus) was transferred to the 96-well plate seeded with Vero-TMPRESS2 cells and incubated at 37°C for 2 h. After 2 h, an additional 40 µL of DMEM supplemented with 20% FBS and 2% PenStrep was added to the cells. After 16-20 h, 40 µL of One-Glo-EX substrate (Promega) was added to each well and incubated on a plate shaker in the dark for 5 min. Relative luciferase units were read using a Biotek plate reader. Relative luciferase units were plotted and normalized in Prism (GraphPad): 100% neutralization being cells lacking pseudovirus and 0% neutralizing being cells containing virus but lacking plasma. Prism (GraphPad) nonlinear regression with “[inhibitor] versus normalized response with a variable slope” was used to determine ID50 values from curve fits with 2-3 repeats. 2-4 biological replicates were carried out for each sample.

### Depletion of SARS-CoV-2 S-binding antibodies from polyclonal plasma

100 µL of Invitrogen His-Tag Dynabeads (ThermoFisher 10104D) were used for prefusion S_2_ depletion, whereas 400 μL was used for S, NTD, and RBD depletion. Vortexed beads were aliquoted into microcentrifuge tubes and incubated on an Invitrogen DynaMag-2 Magnet (ThermoFisher 12-321-D) for two minutes. The supernatant was discarded and beads were washed with 300 µL TBST. After a two-minute incubation on the magnet, the supernatant was discarded and 20 μg of prefusion S_2_ or 80 μg of S, NTD, or RBD in 300 μL TBST was left to incubate with the beads for 1 h at room temperature. The magnet was used and supernatant discarded before being washed with 300 μL TBST. The beads were spun down, put on the magnet, and excess liquid was removed before addition of 20-40 μL of plasma of interest. The plasma was left to incubate with the beads for 1 h at 37 C before being removed. For depletion of postfusion S_2_, 50 µL of Pierce NHS-Activated Magnetic Beads (ThermoFisher 88826) were placed on a magnet. The liquid was removed and beads were washed with 300 µL ice-cold 1 mM hydrochloric acid before addition of 20 µg postfusion S_2_ and incubated for 2 hours at room temperature. The beads were magnetized, washed twice with 300 µL 0.1 M glycine pH 2, and washed once with 300 µL ultrapure water. 300 µL 3 M ethanolamine pH 9.0 was added and the beads were incubated for 2 hours at room temperature before being washed once with ultrapure water. The beads were spun down, put on the magnet, and excess liquid removed before addition of 20 µL of plasma of interest. The plasma was left to incubate with the beads for 1 h at 37 C before being removed.

